# Molecular atlas of postnatal mouse heart development

**DOI:** 10.1101/302802

**Authors:** Virpi Talman, Jaakko Teppo, Päivi Pöhö, Parisa Movahedi, Anu Vaikkinen, S. Tuuli Karhu, Kajetan Trošt, Tommi Suvitaival, Jukka Heikkonen, Tapio Pahikkala, Tapio Kotiaho, Risto Kostiainen, Markku Varjosalo, Heikki Ruskoaho

## Abstract

**Rationale:** Mammals lose the ability to regenerate their hearts within one week after birth. During this regenerative window, cardiac energy metabolism shifts from glycolysis to fatty acid oxidation, and recent evidence suggests that metabolism may participate in controlling cardiomyocyte cell cycle. However, the molecular mechanisms mediating the loss of postnatal cardiac regeneration are not fully understood.

**Objective:** This study aims at providing an integrated resource of mRNA, protein and metabolite changes in the neonatal heart to identify metabolism-related mechanisms associated with the postnatal loss of regenerative capacity.

**Methods and Results:** Mouse ventricular tissue samples taken on postnatal days 1, 4, 9 and 23 (P01, P04, P09 and P23, respectively) were analyzed with RNA sequencing (RNAseq) and global proteomics and metabolomics. Differential expression was observed for 8547 mRNAs and for 1199 of the 2285 quantified proteins. Furthermore, 151 metabolites with significant changes were identified. Gene ontology analysis, KEGG pathway analysis and fuzzy c-means clustering were used to identify biological processes and metabolic pathways either up- or downregulated on all three levels. Among these were branched chain amino acid degradation (upregulated at P23) and production of free saturated and monounsaturated medium- to long-chain fatty acids (upregulated at P04 and P09; downregulated at P23). Moreover, the HMG-CoA synthase (HMGCS)-mediated mevalonate pathway and ketogenesis were transiently activated. Pharmacological inhibition of HMGCS in primary neonatal rat ventricular cardiomyocytes reduced the percentage of BrdU+ cardiomyocytes, providing evidence that the mevalonate and ketogenesis routes may participate in regulating cardiomyocyte cell cycle.

**Conclusions:** This is the first systems-level resource combining data from genome-wide transcriptomics with global quantitative proteomics and untargeted metabolomics analyses of the mouse heart throughout the early postnatal period. This integrated multi-level data of molecular changes associated with the loss of cardiac regeneration may open up new possibilities for the development of regenerative therapies.

## Introduction

Adult human hearts possess a negligible regenerative capacity and therefore cell loss in response to myocardial infarction (MI) leads to scar formation, remodeling of the surrounding myocardium, progressive impairment of cardiac function and eventually to heart failure (reviewed in ^1,2^). Approximately only 0.5-1% of human cardiomyocytes are renewed annually through proliferation of existing cardiomyocytes.^3,4^ This is insufficient for regeneration after MI, which can cause a loss of up to 25% (~1 billion) of cardiomyocytes.^5^ The renewal rate is higher in infants and adolescents than in the elderly, indicating higher regenerative potential in children.^3, 6^ Remarkably, certain amphibians and fish, as well as neonatal rodents can fully regenerate their hearts after an injury.^7-9^ This occurs predominantly through proliferation of remaining cardiomyocytes in the areas adjacent to the injury.^8, 10^ However, rodents lose this regenerative capacity within 7 days after birth due to cardiomyocyte cell cycle withdrawal,^8^ which is considered to represent a major hurdle for the development of regeneration-inducing therapies.^11-13^ In line with cardiac regeneration in neonatal rodents, full functional recovery of a newborn baby with a massive MI suggests that humans possess a similar intrinsic capacity to regenerate their hearts at birth.^14^ It is therefore crucially important to elucidate the molecular mechanisms mediating the postnatal loss of cardiac regeneration.

Upon birth, the resistance of pulmonary circulation drops dramatically due to the expansion of lung alveoli, causing an increase in systemic blood pressure and shunt closure. This in turn induces a steep increase in arterial blood oxygen content. Increased cardiac workload and altered metabolic environment induce a switch from anaerobic glycolysis, which is the main source of energy in the embryonic heart, to mitochondrial fatty acid β-oxidation soon after birth.^15^ This switch is controlled by signaling that involves hypoxia-inducible factor 1α (HIF1α) and HAND1.^16^ Upon birth, HAND1 expression is rapidly down-regulated, allowing cardiomyocytes to shut down glycolysis and initiate lipid oxidation. Failure to downregulate HAND1 and thereby initiate lipid oxidation is fatal, emphasizing that oxidative metabolism is a prerequisite for efficient energy production in the postnatal heart. Moreover, the peroxisome proliferator-activated receptor (PPAR) signaling plays an important role in activating lipid metabolism.^17^

The profound changes in the energy metabolism of cardiomyocytes are associated with alterations in mitochondria: the fetal type mitochondria undergo mitophagy and are replaced with mature adult-type mitochondria in a Parkin-dependent process to allow more efficient ATP production.^18^ Increased oxidative metabolism promotes reactive oxygen species (ROS) production and thereby induces a DNA damage response, which is believed to contribute to cardiomyocyte cell cycle arrest.^19^ However, oxidative DNA damage does not fully correlate with cardiomyocyte cell cycle withdrawal in humans and is thus insufficient to explain the loss of regenerative capacity.^20^ On molecular level, endogenous mechanisms known to regulate cardiomyocyte proliferation include neuregulin-ERBB2 signaling, Hippo-YAP pathway, and the transcription factor GATA4, among others.^11, 21-23^ However, even though metabolic pathways are controlled by the same signals that regulate cell proliferation,^24^ their significance in cardiac regeneration and cardiomyocyte proliferation is yet uncovered.

In search of regenerative therapies, a number of studies have utilized transcriptomics to identify mechanisms regulating cardiac regeneration (for recent examples, see^25-27^). However, proteomic and metabolomic changes have not been thoroughly investigated. Here we report the first integrated study combining genome-wide RNA sequencing (RNASeq), global proteomics and untargeted metabolomics to characterize in detail the metabolic changes occurring during the postnatal heart development.

## Materials & Methods

The experimental design is summarized in Figure 1. Two separate sets of mouse ventricular tissue samples were used. Set 1 encompassed samples from postnatal days 1, 4 and 9 (P01, P04 and P09, respectively) representing hearts with full regenerative capacity (P01), partial regenerative capacity (P04) and negligible regenerative capacity (P09), and was used for proteomics and metabolomics. In order to validate the results, a second set of samples was analyzed. In set 2, an additional sample group from 23 day-old mice (P23) was included to discriminate phenomena related to heart growth. The set 2 samples were subjected to transcriptomics using pooled samples (3 hearts/ sample) and proteomics and metabolomics analyses (no pooling). The methods and bioinformatics analyses are shown in Figure 1B. For transcriptomics analyses, statistically significant differential expression was defined as fold change > 1.5 and q < 0.01. For proteomics, metabolomics and bioinformatics analyses, q < 0.01 with no fold change limit was considered statistically significant.

**Figure 1.**
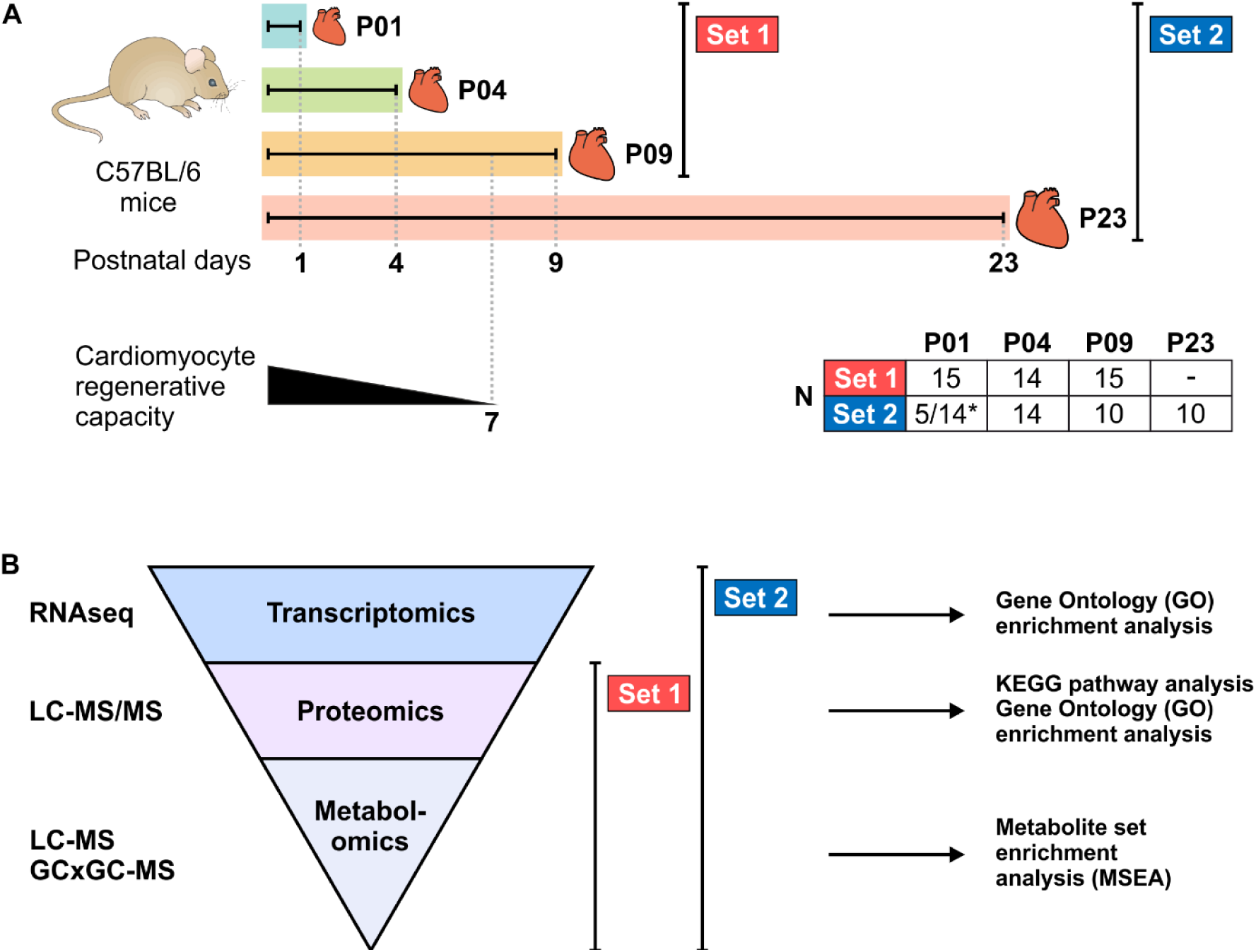
Experimental design for the multi-omics analysis of postnatal mouse hearts. **A**, Two separate sets of mouse ventricular tissue samples collected on postnatal days 1,4, 9, and 23 (P01, P04, P09, and P23, respectively) were used. The postnatal loss of cardiac regenerative capacity is illustrated for comparison, and numbers of animals in each sample group are presented in the table. ^∗^N numbers are indicated for metabolomics (5) / proteomics (14). **B**, Analysis techniques and bioinformatics analyses used in the study. GCxGC-MS, two-dimensional gas chromatography-mass spectrometry; LC-MS/MS, liquid chromatography-tandem mass spectrometry; RNAseq, RNA sequencing.

Detailed Methods are available in the Supplemental Material and detailed protocols from the corresponding authors on reasonable request.

The RNAseq data will be made publicly available at the NCBI gene expression and hybridization array data repository (GEO, http://www.ncbi.nlm.nih.gov/geo/: accession number [to *be added*]). The proteomics raw data will be made available in MassIVE (https://massive.ucsd.edu/ProteoSAFe/static/massive.isp: accession number [to *be added*]. All other omics data are available in the Online Data Supplements (S1-S5).

## Results

### Transcriptomics

In order to analyze postnatal gene expression changes, RNAseq was carried out with ventricular tissue samples using three pooled samples from altogether nine hearts at each time point. Principal component analysis (PCA) of RNAseq data showed clear grouping of samples and separation of sample groups (Supplemental Figure S1). Figure 2A presents hierarchical clustering and a heat-map of 1000 genes with most significant changes between P01 and P04. Altogether 8547 individual protein-coding genes were up- or downregulated statistically significantly (q < 0.01, fold change > 1.5); their numbers for each time point comparison are presented in Figure 2B. The greatest changes in gene expression were observed between time point comparisons P01 to P23 and P04 to P23. The top 10 up- and downregulated genes are presented in Supplemental Figure S2. The expression levels of cardiomyocyte-specific structural proteins exhibited an anticipated pattern with a switch from *Myh7* to *Myh6* and from *Tnni1* to *Tnni3* within the early postnatal period (Figure 2C).^28, 29^ Also the expression profiles of the main cardiomyocyte ion channels (Supplemental Figure S3A) were in line with previous reports.^30^ The expression patterns of control genes are presented in Supplemental Figure S3B. Expression of *Actb*, *Rpl4*, *Rpl32*, *Tbp*, *Oaz1* and *Pgk1* did not change significantly, whereas several other genes either generally used or recommended as control genes^31^, such as *Gapdh*, were up- or downregulated (q < 0.01 and fold change > 1.5) in at least one time-point comparison. Differential expression analysis of all protein-coding genes is available in Supplemental Dataset S1.

**Figure 2.**
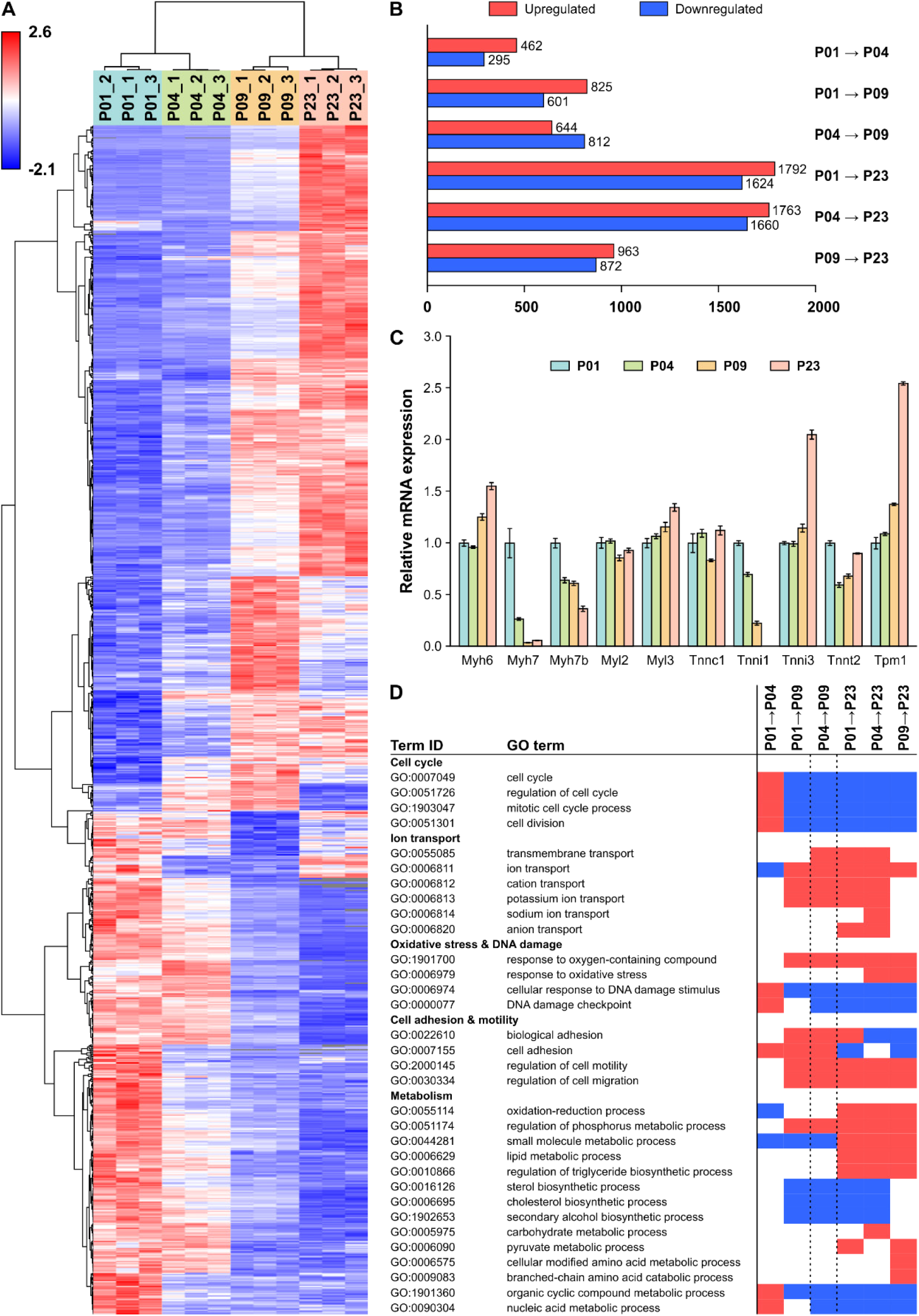
Gene expression changes in the neonatal mouse heart. **A**, Heatmap of the top 1000 genes with smallest q values between P01 and P04**. B**, The numbers of up- and downregulated genes (q < 0.01, fold change > 1.5). C, Expression patterns of selected cardiomyocyte-specific structural proteins. The data were normalized to P01 and are expressed as mean±SEM (n=3 pooled samples, each from 3 hearts). D, Selected significantly enriched (q < 0.01) biological process GO terms for each time point comparison from GSEA. Blue indicates downregulation and red indicates upregulation. All gene symbol explanations are available in the Supplemental Dataset S1.

To investigate biological processes linked to the observed gene expression changes we carried out gene ontology (GO) enrichment analysis and changes with q < 0.01 were considered significant. Changes in biological processes linked to cell proliferation, cardiac muscle, cell adhesion and motility, ion transport and development of immune response were highlighted among the enriched GO terms (Figure 2D and Supplemental Dataset S2). In accordance with increased oxidative metabolism, genes linked to oxidative stress were upregulated at P23 compared to P04 or P09. DNA damage response was upregulated from P01 to P04, whereafter it was downregulated in all other comparisons. Of the GO terms associated with cellular metabolism, oxidation-reduction processes were upregulated at P23 compared to P01, P04 and P09, carbohydrate metabolism was upregulated from P04 to P23 and pyruvate metabolism from P01 to P23 and P09 to P23. Sterol biosynthesis was downregulated in all other time point comparisons except P01 to P04 and P09 to P23, indicating that the transcript-level changes take place mainly between postnatal days 4 and 9. Amino acid metabolism and branched chain amino acid (BCAA) catabolism were upregulated from P09 to P23.

### Proteomics

To quantify protein abundances in ventricular tissue, we used a shotgun proteomics approach. On average, 1140 and 1337 distinct proteins were quantified in individual samples in sample sets 1 and 2, respectively (Figure 3A). The numbers and fold changes of differentially expressed proteins are shown in Supplemental Figure S4A (set 1) and Figure 3B (set 2). The corresponding lists of differentially expressed proteins, as well as all quantified proteins and their label-free quantification intensities are presented in Supplemental Dataset S3. Hierarchical clustering of the data shows that the sample groups were separated from each other in both sample sets (Figure 3C and Supplemental Figure S4B), which was also seen in PCA (Supplemental Figure S5). GO enrichment analysis and KEGG pathway analysis were used to identify up- and downregulated processes and activated/ inactivated pathways (Supplemental Dataset S3). Oxidative metabolism and lipid metabolism were highlighted as upregulated phenomena at P04, P09 and P23 compared to P01, while nucleic acid metabolism and glycolysis were highlighted in the downregulated gene ontologies from P01 to P04, peptide metabolism and protein synthesis from P01 to P09 and RNA processing and peptide metabolism from P01 to P23.

**Figure 3.**
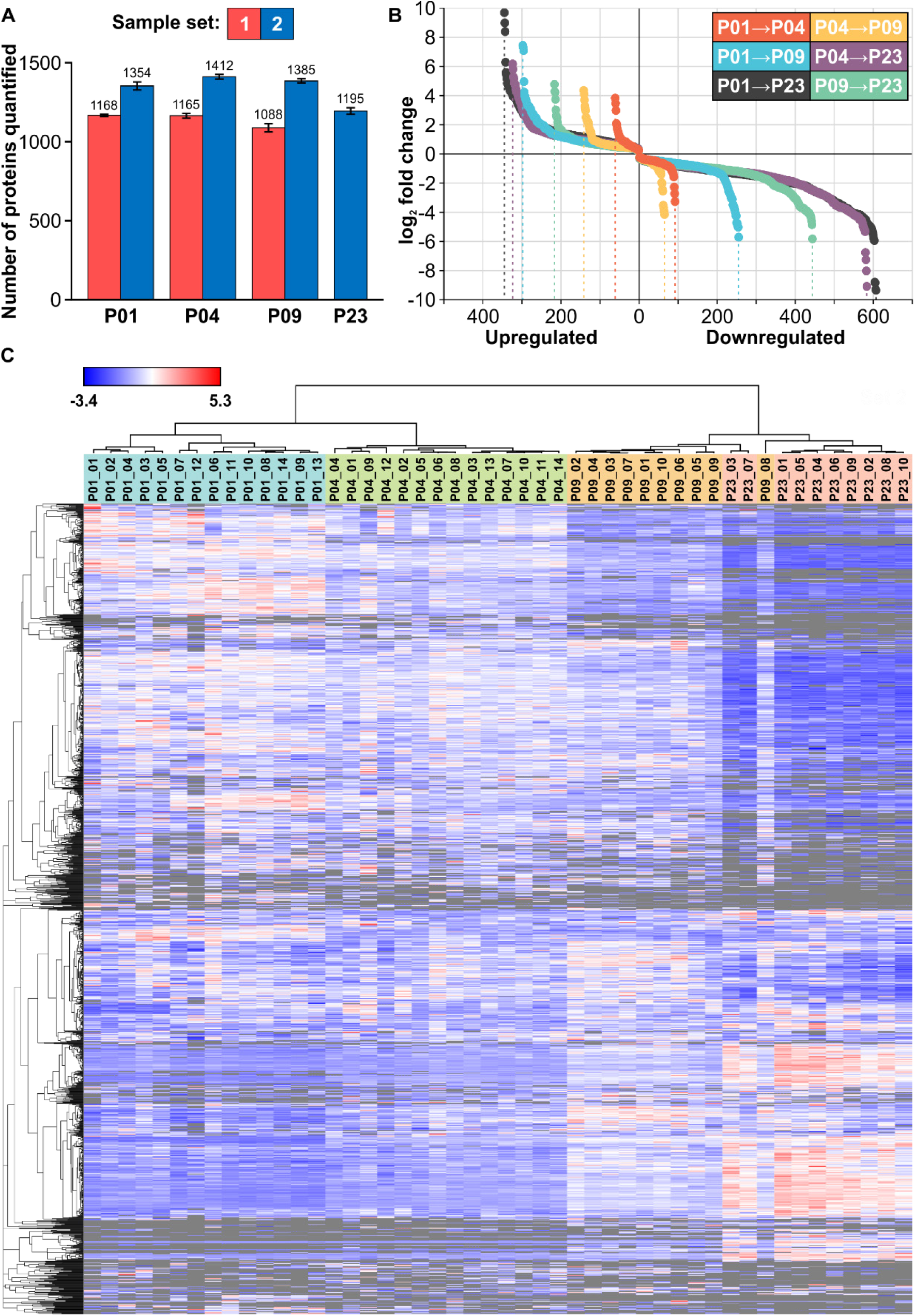
Proteomic changes in the neonatal mouse heart. **A**, The number of proteins quantified in each sample group, expressed as mean±SEM (identification FDR < 0.01 on both peptide and protein level). **B**, The numbers and fold changes of differentially expressed (q < 0.01) proteins in sample group comparisons**. C**, Hierarchical clustering of proteins and samples. All proteins detected in > 2/3 samples of at least one sample group are included in the heatmap. Gray color indicates missing values (protein not detected). **B-C**, Data presented is from set 2; corresponding images for set 1 are in Supplemental Figure S4.

The postnatal increased activity of energy metabolism as well as the shift from carbohydrates to fatty acids as the primary source of ATP were well represented in the proteomics data. Several components (ENO1, PFKL, PGK1) of the HIF-1 signaling pathway that promotes anaerobic metabolism in the embryonic heart were down-regulated after birth (Supplemental Table 1). Furthermore, many key proteins related to fatty acid metabolism, degradation and oxidative phosphorylation were upregulated after P01. This was paralleled with upregulation of most of the detected components of PPAR signaling, which promotes fatty acid oxidation, after P01. In line with the increasing mitochondrial content and maturation in cardiomyocytes, most mitochondrial proteins were highly upregulated from P01 to P23. These included not only enzymes linked to oxidative metabolism but also structural (IMMT) and regulatory (PHB, DNAJA3) proteins (Supplemental Table 1). Furthermore, the mitochondrial isoforms of superoxide dismutase (SOD2) and peroxiredoxin (PRDX3, PRDX5) were strongly upregulated at P09 and P23 (Supplemental Table 1), reflecting a response to oxidative stress.

Analysis of key proteins of cell cycle regulation and signaling pathway activation with proteomics is challenging due to low abundance of these proteins. Nevertheless, expression of proteins related to microtubule dynamics during mitosis (DYNC1H1, PAFAH1B1, RHOA, and BUB3) decreased with increasing postnatal age (Supplemental Table 1). Most of the detected 14-3-3 proteins (YWHAB, YWHAE, YWHAH, YWHAQ, and YWHAZ), some of which regulate cell cycle and cardiomyocyte proliferation,^32, 33^ were constantly downregulated from P01 to P23 in set 2 samples (Supplemental Table 1). Components of the canonical Wnt pathway were also detected (Supplemental Table 1), and the decreased abundance of β-catenin (CTNNB1) and its nuclear interaction partner Pontin52 (RUVBL)^34^ indicate attenuation of canonical Wnt pathway activity from P01 to P09.

### Metabolomics

For the untargeted metabolomics analysis, we used two complementary techniques to increase the metabolite coverage, namely liquid chromatography-mass spectrometry (LC-MS) and two-dimensional gas chromatography coupled to mass spectrometry (GCxGC-MS), and combined the data from the two sample sets using linear mixed effect (LME) model. Quality control results are available in the Supplemental Material. In total, 805 and 162 metabolic features changed statistically significantly (q < 0.01) in at least one of the group comparisons (P01→P04, P01→P09, P01→P23) for LC-MS and GCxGC-MS analyses, respectively (Supplemental Dataset S4). Based on the MS/MS analysis of metabolic features, 151 significantly up- or downregulated metabolites were identified and are shown in Figure 4.

**Figure 4.**
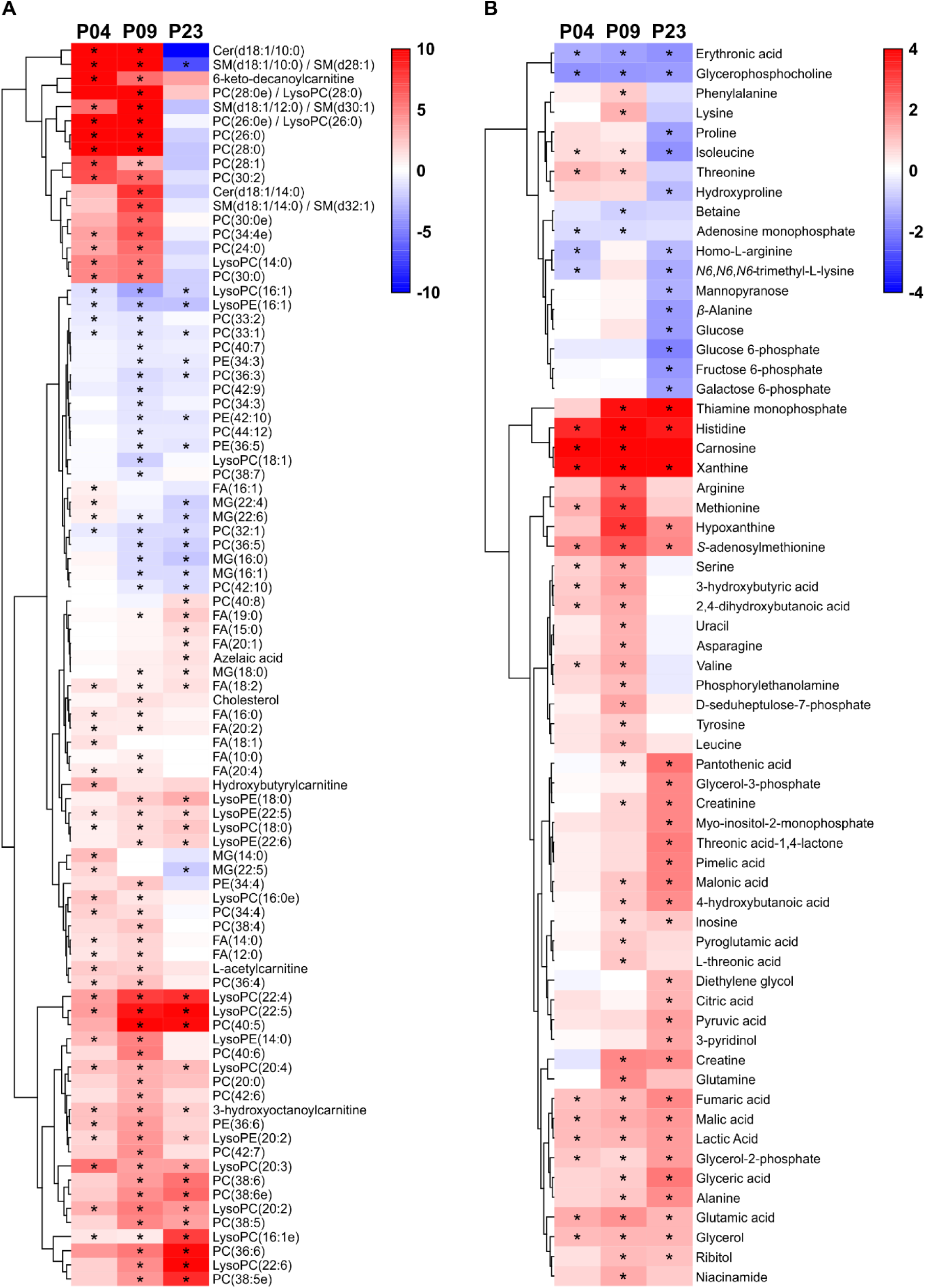
Metabolite changes in the postnatal mouse heart. **A**, Heatmap of linear mixed effect (LME) model estimates for lipids. **B**, Heatmap of LME model estimates for polar metabolites with Glass’ delta effect sizes from P01. ^∗^q < 0.01 compared to P01. Cer, ceramide; SM, sphingomyelin; PC, phosphatidylcholine; PE, phosphatidyle-thanolamine; LysoPE, lysophosphatidylethanolamine; LysoPC, lysophosphatidylcholine; MG, monoacylglycerol; FA, fatty acid.

In line with the transcriptomics and proteomics results, the metabolomics data shows a distinct transition from carbohydrates to lipids as the main source of energy. The abundances of glucose and sugar derivatives (glucose, glucose-6-phosphate, fructose-6-phosphate, galactose-6-phosphate) were downregulated at P23 compared to P01 (Figure 4B), while the abundances of most fatty acids and components of glycerolipid metabolism (glycerol-3-phosphate, glycerol-2-phosphate, glycerol, glyceric acid) increased after P01 (Figure 4A). The increased fatty acid β-oxidation was also reflected as increased abundance of acylcarnitines at P04 and P09 (Figure 4A). However, most metabolites of the citric acid cycle (citric acid, fumaric acid, pyruvic acid, malic acid, lactic acid), pentose-phosphate pathway (D-seduloheptulose-7-phosphate) and glycolysis (lactic acid, pyruvic acid) exhibited a constant rise with increasing postnatal age (Figure 4B), reflecting a total increase in energy metabolism. No changes were detected in the levels of citric acid cycle metabolites succinic acid and α-ketoglutarate (Supplemental Dataset S4). Overall, these data indicate a transition from carbohydrates to fatty acids as the main source of energy and a total increase in energy metabolism over the early postnatal period.

In addition to the general increase in fatty acid abundance, various lipid species (phospholipids, lysophospholipids, monoacylglycerides and fatty acids) displayed interesting changes in abundance (Figure 4A). For phospholipids, there was a general trend towards increased saturation level along with increasing postnatal age. All phospholipids showing a constant decrease from P01 to P23 were unsaturated, and many of them polyunsaturated. The phospholipid species with saturated medium-to-long-chain fatty acids exhibited an increase at P04 and P09 followed by a decrease at P23. Several lysolipid species (LysoPC and LysoPE) increased at P04 and remained constant or increased further through P09 to P23, with the exception of lysolipids with fatty acids 16:1 and 18:1, which decreased after P01, or 14:0, which decreased after P04. Interestingly, myristic acid (fatty acid 14:0) exhibited a comparable pattern both as a free fatty acid and when incorporated into other lipid species (such as ceramide, sphingomyelin, or phosphocholine): the abundance increased at P04 and P09 and decreased at P23. A similar pattern was also observed for other medium-chain saturated fatty acid species (Figure 4A).

The levels of most amino acids displayed an initial increase at P04 and/or P09 followed by a decrease at P23 (Figure 4B). In contrast to other (proteinogenic) amino acids, the abundance of glutamic acid, alanine and histidine increased initially, but remained significantly higher at P23 compared to P01 (Figure 4B). The increase in amino acid abundance from P01 to P09 likely reflects the active protein synthesis required during cardiomyocyte growth and maturation.

We also observed significant changes in purine metabolism, in particular among the metabolites of adenosine monophosphate (AMP) catabolism. The levels of AMP decreased already at P04, which was paralleled by an increase in the abundance of its degradation pathway metabolites inosine, hypoxanthine and xanthine (Figure 4B). This likely reflects the alterations in the energy metabolism, as high AMP abundance immediately after birth would promote fatty acid β-oxidation through the AMP activated protein kinase (AMPK).^35^ In line with the postnatal increase in cardiac workload, we also observed increased abundance of creatine and creatinine at P09 and P23 when compared to P01 (Figure 4B). Other interesting metabolite findings include the increase of the ascorbic acid metabolites L-threonic acid and threonic acid 1,4-lactone on P09 and P23, respectively; reflecting increased oxidative stress with increasing postnatal age.

### Multi-omics integration

For comprehensive multi-omics integration, we utilized fuzzy c-means clustering, in which the transcripts, proteins, and metabolites were assigned into one or more clusters based on their abundance patterns (Figure 5A, Supplemental Figure S6). We compared transcriptomics and proteomics clusters on transcript/protein levels, but also for enriched GO terms and KEGG pathways (q < 0.01). The percentage of proteomics clusters covered by the RNAseq clusters are shown in Figure 5B for the proteins/transcripts and biological process GO terms, and in Supplemental Figure S7 for cellular component and molecular function GO terms and KEGG pathways. Selected enriched GO terms and KEGG pathways and their enrichment in each transcriptomics and proteomics cluster are presented in Figure 5C (cellular component GO terms in Supplemental Figure S7).

**Figure 5.**
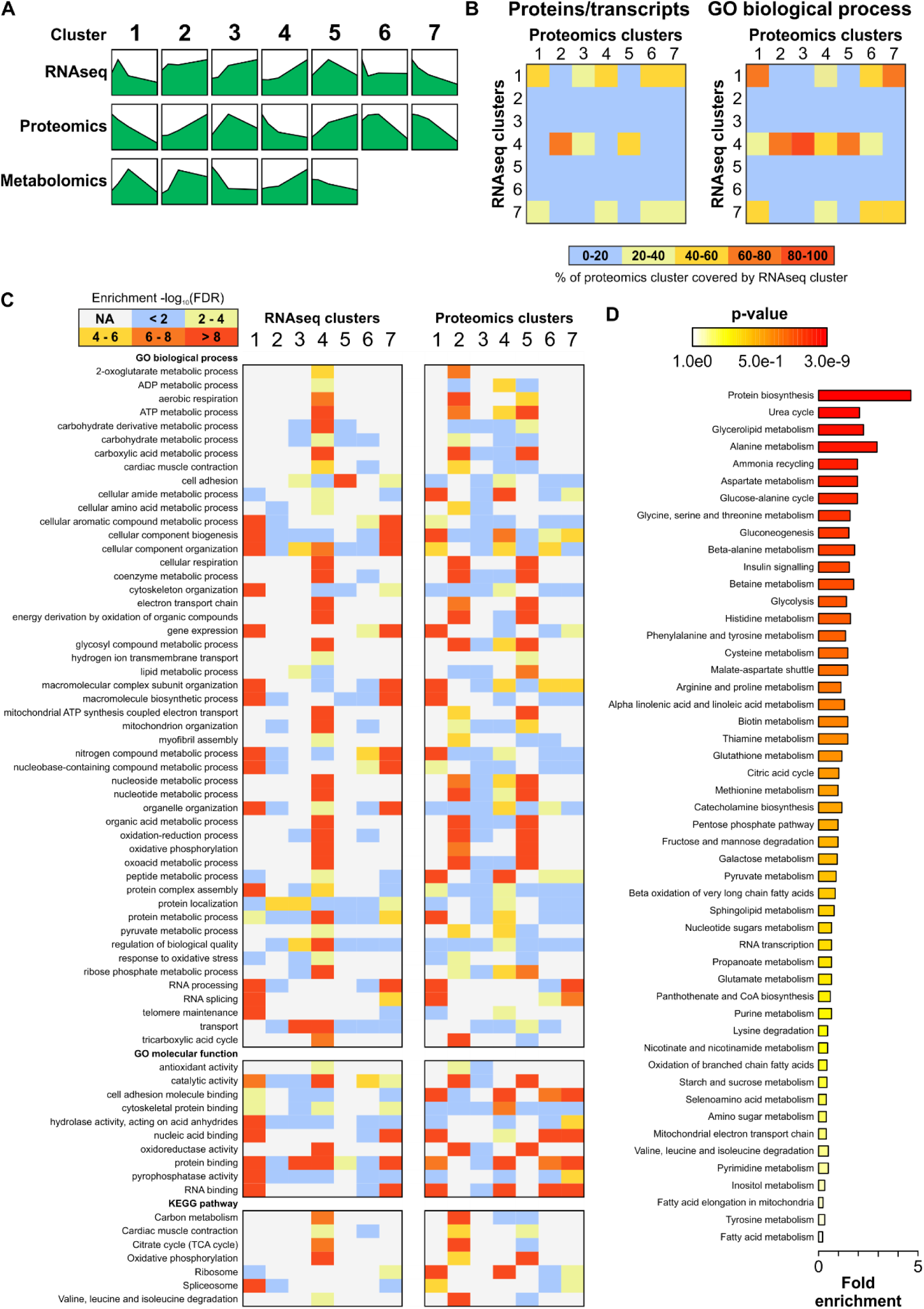
Multi-omics integration with fuzzy c-means clustering. **A**, The median abundance patterns of mRNAs, proteins, and metabolites in each cluster. Detailed images of clusters and the scales for the Y-axis are in Supplemental Figure S6. **B**, The percentage of proteomics clusters covered by the RNAseq clusters at the level of transcripts/proteins (left) and enriched biological process GO terms (q < 0.01) in the clusters (right). **C**, A heatmap of -log_10_ FDR-values of selected GO terms and KEGG pathways in RNAseq and proteomics clusters. **D**, Metabolite set enrichment analysis performed on the union of metabolites in all clusters indicating the biological processes associated with metabolite changes.

Since fuzzy clustering could only be carried out separately for the two sample sets, it provided only little more insight to the metabolomics data compared to the LME model. Therefore, instead of analyzing the individual metabolomics clusters, we performed metabolite set enrichment analysis (MSEA) to all significantly changed and identified metabolites in order to investigate, which of the metabolic pathways were consistently changed. Based on MSEA, the three most significantly enriched metabolic pathways were protein biosynthesis, urea cycle and glycerolipid metabolism (Figure 5D). Enrichment of multiple individual amino acid metabolic pathways was also observed. The high enrichment of urea cycle metabolites correlated with the changes in individual metabolites, such as arginine and fumarate (Figure 4B). Not all identified urea cycle metabolites however exhibited statistically significant changes (such as aspartate and urea; Supplemental Dataset S4).

The KEGG pathway ‘glycolysis and gluconeogenesis’ is presented in the Supplemental Figure S8 as an example. In line with the data from the individual omics analyses, glycolysis was not uniformly activated or inactivated. Instead, the genes and proteins in the beginning and the end of the pathway were either upregulated at later time points or exhibited variable abundance patterns, while the genes and proteins in the middle of the pathway were either downregulated with increasing postnatal age or displayed variable expression patterns. Most enzymes of the related pyruvate metabolism pathway were also differentially expressed on mRNA and/or protein level within the early postnatal period (Supplemental Figure S9).

### Branched Chain Amino Acid Catabolism

To gain deeper insight into the significant enrichment of the KEGG pathway ‘valine, leucine and isoleucine degradation’ in transcriptomics and proteomics clusters showing upregulation throughout the postnatal period, we focused on BCAA catabolism in more detail. The concentrations of valine, leucine and isoleucine increased from P01 to P09, whereafter they decreased to P01 levels or below at P23 (Figure 6A). Most enzymes in the BCAA degradation pathway were significantly upregulated at P23 compared to P01 on either transcript or protein levels, or both (Figure 6B). On the mRNA level, most differentially expressed genes were upregulated between P09 and P23, thus correlating directly with the BCAA concentrations. Of the 37 quantified individual proteins on this pathway, 19 and 31 proteins were upregulated at P09 and P23, respectively, when compared to P01 (Set 2 samples; Supplemental Dataset S3). The rate-limiting step of BCAA catabolism is mediated by the branched-chain α-ketoacid dehydrogenase (BCKDC or BCKDH) complex, whose activity is controlled by inactivating phosphorylation and activating dephosphorylation. Both the α-subunit of the complex, *Bckdha*, and the protein phosphatase responsible for BCKDH activation (*Ppm1k*) were significantly upregulated on the mRNA level at P23, and BCKDHA protein abundance was upregulated at P23 (PPM1K was not detected; Supplemental Table 1).

**Figure 6.**
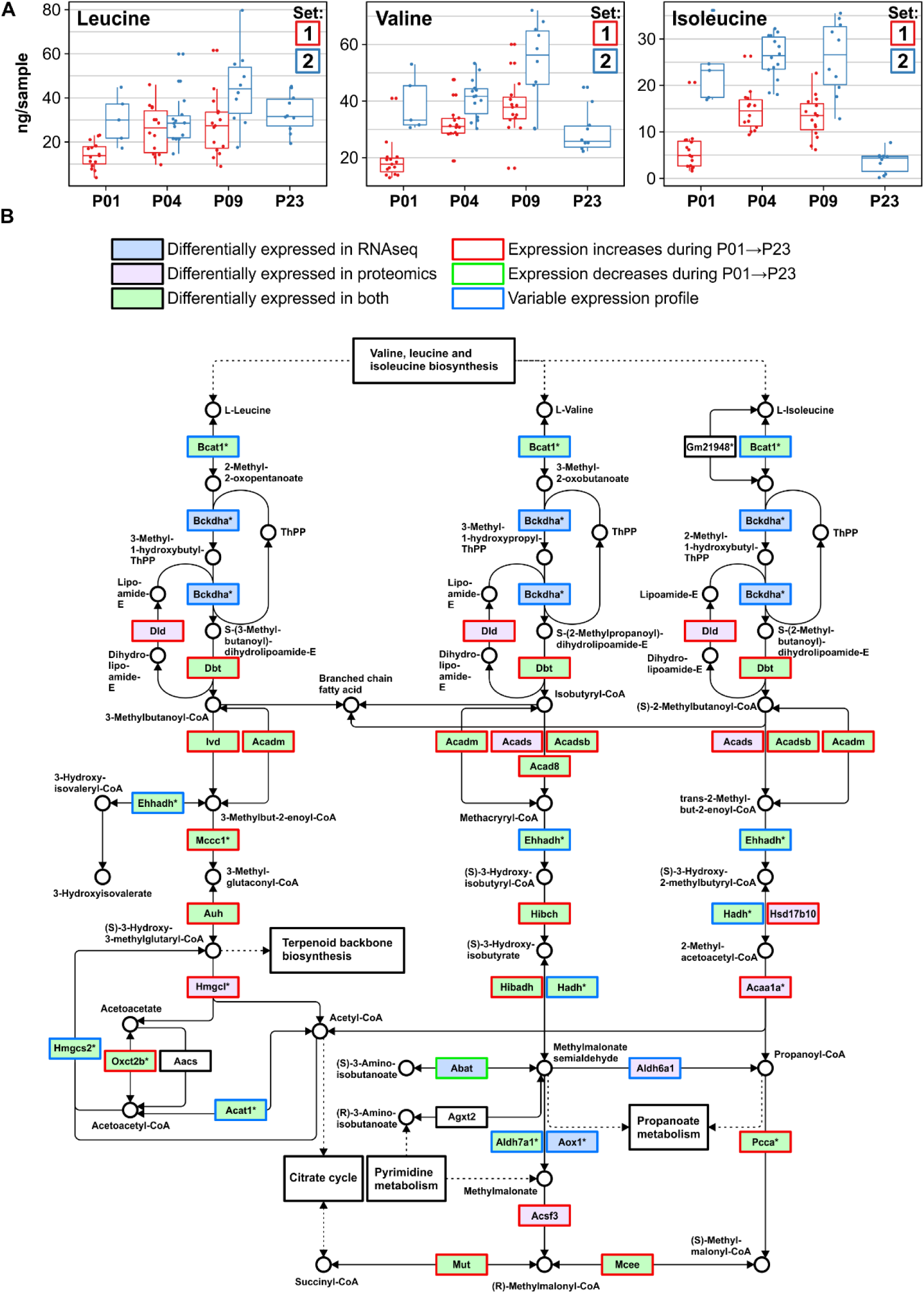
Postnatal changes in branched chain amino acid degradation in the mouse heart. **A**, Concentrations of branched chain amino acids valine, leucine, and isoleucine in mouse ventricular tissue. **B**, The KEGG pathway map of valine, leucine and isoleucine degradation indicating up- and downregulated transcripts and proteins. ^∗^ Indicates that multiple enzymes may catalyze the same reaction. All gene symbol explanations are available in the Supplemental Dataset S1. The KEGG pathway image is modified and reprinted from http://www.genome.jp/kegg/ with permission from the Kyoto Encyclopedia of Genes and Genomes.

### Fatty acid metabolism

The main enzymes regulating the abundance of free fatty acids are the fatty acid synthase (FASN); long-chain acyl-CoA synthetases (ACSLs), which attach Coenzyme A (CoA) to free fatty acids; and acyl-CoA thioesterases (ACOTs), which remove the CoA from acyl-CoA thus releasing free fatty acid (Figure 7A). The relative mRNA levels of FASN, ACOTs and ACSLs are presented in Figure 7B. Downregulation of *Acot1*, *Acot2*, *Acsl3*, *Acsl4* and *Fasn* was observed after P01, whereas *Acot13* was upregulated at P23. In general, there was a relatively poor correlation between protein (Figure 7C) and mRNA expression. The abundance of ACOT isoforms increased, except for ACOT1, which was first upregulated at P04 and P09, followed by downregulation at P23. The abundance of ACSL1 increased over the postnatal period. FASN, however, was strongly downregulated at P23 to undetectable levels, correlating with the mRNA expression pattern. The products of the cytosolic ACOT1 and mitochondrial ACOT2, saturated and monounsaturated medium-to-long-chain fatty acids,^36^ exhibited almost identical patterns with an initial increase peaking at P04 or P09, followed by strongly diminished levels at P23 (Figure 7D) and thus correlating with ACOT1 abundance. Furthermore, most enzymes in the KEGG pathway ‘fatty acid degradation’ were differentially expressed on mRNA and/or protein level (Supplemental Figure S10). These data reflect the complex regulation of fatty acid levels in the postnatal heart in response to the increased fatty acid abundance from nutrients.

**Figure 7.**
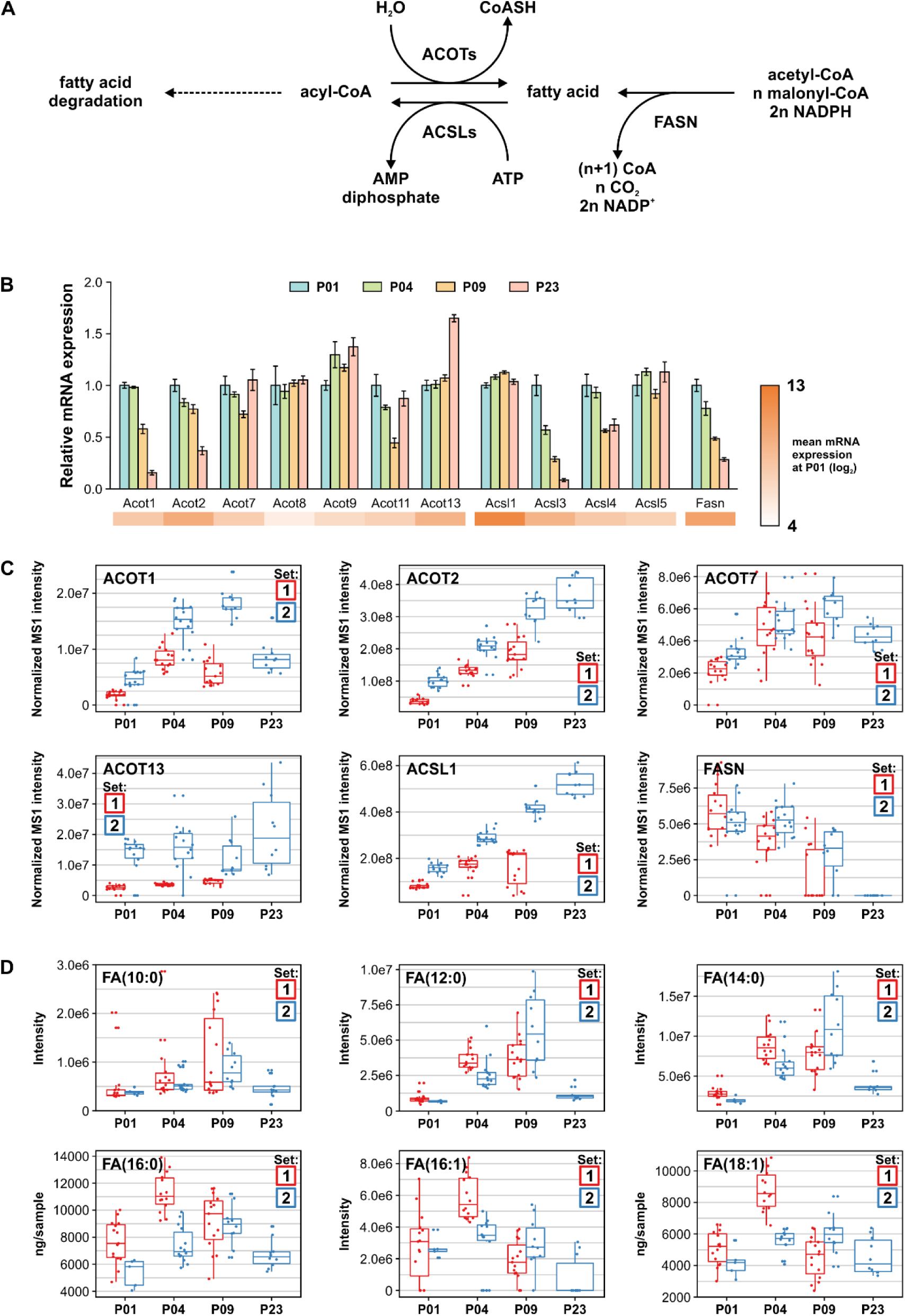
Postnatal changes in fatty acid metabolism in the mouse heart. **A**, The levels of free fatty acids (FAs) are regulated by acyl-coenzyme A thioesterases (ACOTs) that hydrolyze acyl-CoAs to CoASH and free FAs, long-chain-fatty-acid--CoA ligases (ACSLs) that activate free FAs by ligation of CoA, and FA synthase (FASN). **B**, Relative mRNA expression of enzymes regulating the concentrations of free FAs, shown as mean±SEM (n = 3 pooled samples, each from 3 hearts). C, Normalized label-free quantification intensities of FA-regulating enzymes detected in proteomics. D, The abundances or concentrations of selected free FAs.

### The mevalonate pathway and ketogenesis

Since hydroxymethylglutaryl-CoA (HMG-CoA) synthase 2 (HMGCS2) was the most significantly upregulated protein from P01 to P04 and cholesterol biosynthesis was one of the enriched metabolism-related biological processes in GO enrichment analysis, the HMGCS-mediated mevalonate pathway and ketogenesis were investigated in more detail. The HMGCS-catalyzed synthesis of HMG-CoA serves as a substrate for HMG-CoA reductase (HMGCR) in the mevalonate pathway and HMG-CoA lyase (HMGCL) in ketogenesis (Figure 8A). Both the cytosolic *Hmgcs1* and the mitochondrial *Hmgc2* isoforms were downregulated on mRNA level at P09 and P23 compared to P01, whereas there was no change in the mRNA levels of the upstream enzyme acetyl-CoA acetyl-transferase (ACAT1; Figure 8B). Of the enzymes regulating ketone body abundance, expression of *Bdh1* was significantly lower at P09 compared to P01 or P23 and the expression of *Oxct1*, which oxidizes ketone bodies, was upregulated after P09. In the mevalonate pathway, *Hmgcr* expression was downregulated at P09 when compared to P01 or P04 and at P23 when compared to P04. Similarly, mevalonate kinase (*Mvk*), phosphomevalonate kinase (*Pvmk*) and isopentenyl-diphosphate delta isomerase 1 (*Idi1*) were downregulated with increasing postnatal age. Of these enzymes, ACAT1, HMCSS2, HMGCL and OXCT1 were reliably detected in proteomics (Figure 8C). The abundances of ACAT1 and HMGCL increased throughout the early postnatal period, while HMGCS2 peaked at P04 and was downregulated to undetectable levels by P23 and OXCT1 was upregulated from P09 to P23. On the metabolite level, the end product of ketogenesis, 3-hydroxybutyric acid (β-hydroxybutyrate), peaked at P09 and was downregulated at P23 (Figure 8D), correlating with changes in the abundance of HMGCS2, which is the rate-limiting enzyme of ketogenesis, and OXCT1. Metabolites of the mevalonate pathway were not detected or identified; however, the concentrations of cholesterol, which is produced from mevalonate, increased at P09 and decreased thereafter (Figure 8D). Collectively, these results show that the HMGCS-mediated mevalonate pathway and ketogenesis are activated transiently after birth in the mouse heart.

**Figure 8.**
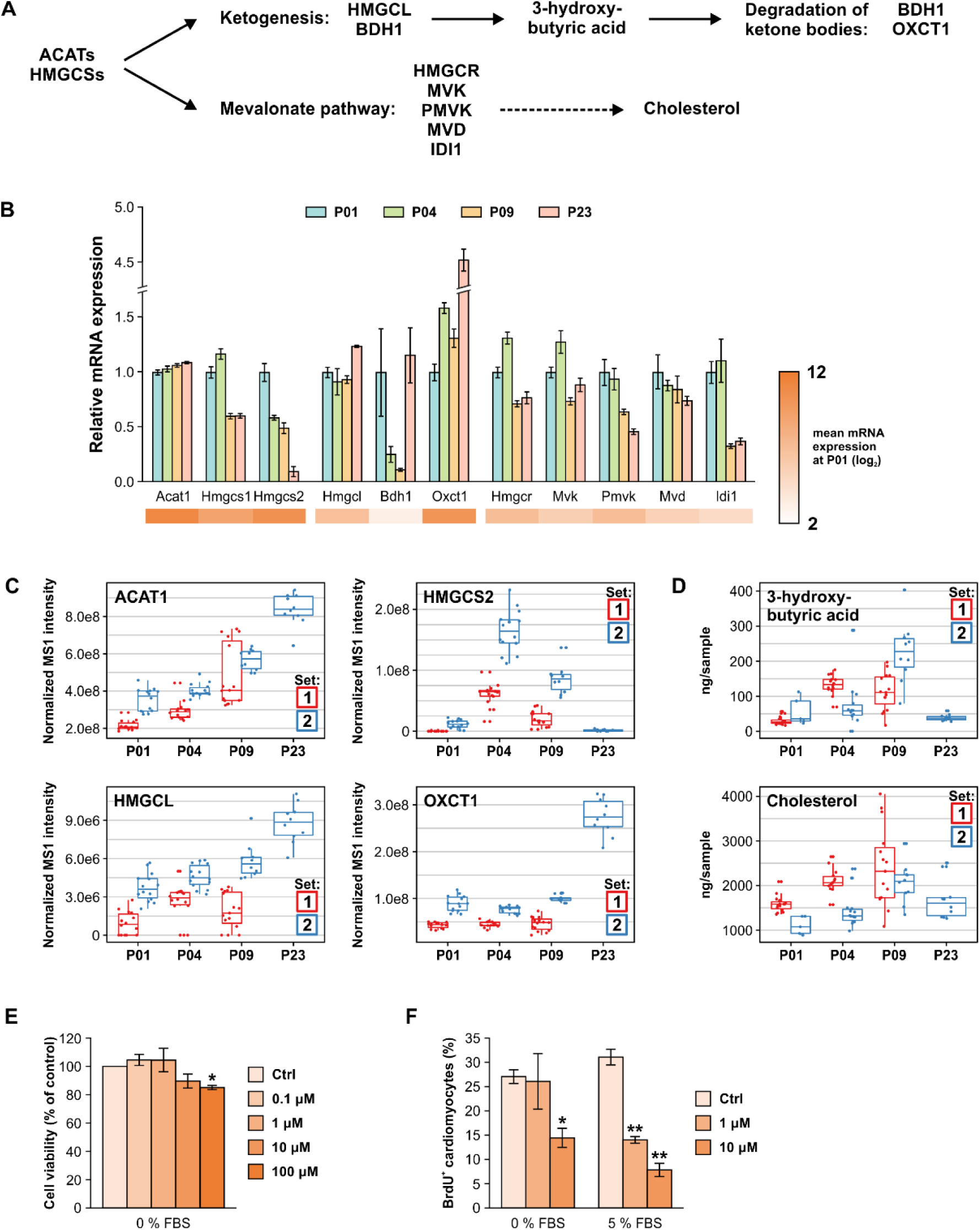
Ketogenesis and mevalonate pathways in the mouse heart. **A**, Acetyl-CoA acetyltransferases (ACATs) and hydroxymethylglutaryl-CoA synthases (HMGCSs) catalyze hydroxymethylglutaryl-CoA (HMG-CoA) synthesis. HMG-CoA serves as a substrate for the ketogenesis route producing 3-hydroxybutyrate and the mevalonate pathway producing mevalonate, which can be further used for cholesterol synthesis. **B**, Relative mRNA expression of selected ketogenesis and mevalonate pathway components, shown as mean±SEM (n = 3 pooled samples, each from 3 hearts). **C**, Normalized label-free quantification intensities of proteins detected in the ketogenesis and mevalonate pathways. **D**, Concentrations of 3-hydroxybutyrate and cholesterol over the early postnatal period. **E-F**, Effect of HMGCS inhibition with hymeglusin on neonatal rat ventricular cardiomyocyte viability **(E)** and proliferation **(F).** Cell viability was assessed using the MTT assay and cell proliferation was quantified as the percentage of BrdU+ cells, bith after a 24-h exposure. The data are expressed as mean±SEM from 3 independent experiments. ^∗^ p < 0.05, ^∗∗^ p < 0.01 compared to control (Ctrl); Welch ANOVA followed by Games-Howell. FBS, fetal bovine serum. All gene symbol explanations are available in the Supplemental Dataset S1.

To evaluate the role of mevalonate pathway and ketogenesis in cardiomyocyte proliferation, we investigated the effects of pharmacological HMGCS inhibition on the viability and proliferation of neonatal rat ventricular cardiomyocytes. The HMGCS inhibitor hymeglusin^37^ induced a ~15% decrease in cell viability at 100 μM but had no effect at lower concentrations (Figure 8E). However, hymeglusin decreased in the percentage of BrdU-positive cardiomyocytes in both serum-free and serum-stimulated conditions already at smaller, non-toxic concentrations (at 10 μM and at 1-10 μM, respectively; Figure 8E). These results indicate that the HMGCS-mediated mevalonate pathway and/or ketogenesis may participate in regulating cardiomyocyte cell cycle activity.

## Discussion

Therapeutic strategies to promote regeneration of the adult human hearts are urgently sought after.^12, 13^ More detailed understanding of the metabolic changes and signaling pathways mediating cardiomyocyte maturation and cell cycle withdrawal is however required for the development of regenerative therapies. In the present work, we employed an integrated multi-omics approach to investigate the metabolic changes occurring in the mouse heart within the early postnatal period in order to identify metabolic pathways associated with the postnatal loss of regenerative capacity. To our knowledge, this is the first study incorporating genome-wide gene expression analysis to global quantitative proteomics and untargeted metabolomics and thus provides an important resource of molecule abundances in the neonatal mouse heart. Furthermore, we highlighted examples of metabolic pathways that exhibited correlative changes on all three levels.

According to previous reports and the present data, the cardiac energy metabolism changes drastically after birth in response to altered nutrient availability and transition to oxygen-rich environment. Increased oxidative metabolism gives rise to ROS causing oxidative stress and DNA damage, which is thought to contribute to cardiomyocyte cell cycle withdrawal.^19, 38^ Remarkably, exposure of adult mice to chronic hypoxia reduces oxidative metabolism and DNA damage and promotes cardiac regeneration after myocardial infarction.^39^ Oxidative DNA damage does not however correlate directly with cardiomyocyte cell cycle withdrawal in humans.^20^ The continued proliferative phenotype of human pluripotent stem cell (hPSC)-derived cardiomyocytes – despite the normoxic environment – also indicates that other mechanisms contribute to the irreversible cell cycle withdrawal. Accordingly, the metabolic switch from glycolysis to fatty acid oxidation, achieved by increased palmitic acid availability and insulin depletion, was recently reported to be sufficient for inducing irreversible cell cycle exit of hPSC-derived cardiomyocytes.^40^ Furthermore, increased fatty acid abundance has been shown to mediate postprandial physiological cardiac hypertrophy in Burmese python.^41^ Administration of a combination of myristic, palmitic and palmitoleic acid to mice or pythons also induces cardiac hypertrophy without pathological fibrosis or activation of the fetal gene program. Here we showed a temporally regulated postnatal increase in the abundance of saturated and monounsaturated medium-chained fatty acids – both as free fatty acids and in various lipid species such as ceramides, sphingomyelins and phosphocholines. This transient increase could serve as the physiological mechanism regulating non-pathological cardiomyocyte hypertrophy during postnatal heart growth, as reported for the Burmese python.^41^ It is also tempting to speculate that the same fatty acids may play a role in driving postnatal cardiomyocyte cell cycle withdrawal as reported for hPSC-derived cardiomyocytes.^40^

Another key finding of this work, not previously described in the context of postnatal heart development, is the temporal regulation of mevalonate pathway, which was strongly activated immediately after birth and downregulated after the regenerative window. The mevalonate pathway plays a role in cancer cell proliferation and is upregulated by several oncogenic signaling routes.^42^ Furthermore, its downregulation has been linked to increased cell size in vivo,^43^ and inhibition of HMGCR, the rate-limiting enzyme of mevalonate pathway, attenuates cell proliferation and increases cell size in various cell types in vitro.^44^ It can therefore be speculated that the temporal activation of mevalonate pathway in the postnatal mouse heart may participate in regulating cardiomyocyte size and proliferation.

Parallel to the mevalonate pathway, we observed a transient postnatal activation of ketogenesis in the mouse heart. The circulating levels of β-hydroxybutyrate, which is mainly produced in the liver, increase temporarily after birth and provide an important source of energy for the developing brain.^45^ However, the observed strong temporal upregulation of HMGCS2, the rate-limiting enzyme of ketogenesis, indicates that ketone body synthesis is also locally regulated and postnatally activated in the myocardium. In addition to its role as a circulating energy source, β-hydroxybutyrate participates in cellular signaling by acting on cell membrane receptors and by directly inhibiting histone deacetylases.^46^ Increased abundance of ketogenic enzymes has been linked to aggressiveness of prostate cancer,^47^ indicating that augmented ketogenesis may provide a proliferative advantage in certain conditions. We further showed that simultaneous inhibition of mevalonate pathway and ketogenesis attenuates neonatal cardiomyocyte proliferation *in vitro.* The transient postnatal upregulation of ketogenesis in the postnatal mouse heart may thus participate in regulating signaling, gene expression and cell cycle in cardiomyocytes.

Unlike other amino acids, the BCAAs valine, leucine and isoleucine are mainly metabolized in other organs than the liver, such as in skeletal and cardiac muscle. They provide nitrogen for maintaining glutamate, alanine and glutamine pools and function as signaling molecules activating mTOR signaling, which regulates cardiac homeostasis and plays a crucial role in cardiac pathophysiology.^48, 49^ In addition, isoleucine has also been reported to inhibit the transport and utilization of fatty acids in skeletal muscle in mice.^50^ The observed increase in BCAA degradation in the postnatal heart is thus in line with the decreased protein synthesis at P23 and the increased in fatty acid metabolism after P01. Even though the abundance of BCAAs is altered in experimental models of pressure overload and myocardial infarction,^51^ the postnatal changes observed here are unlikely to contribute to the postnatal loss of cardiac regeneration, as they predominantly take place after the regenerative window.

Due to methodological limitations, proteomic and metabolomic analyses can only detect a fraction of proteins and metabolites. By using shotgun proteomics, we were able to perform relative quantification of >2000 proteins. For example, hydrophobic (e.g. transmembrane) proteins and proteins with very low abundance were often below detection. To increase metabolite coverage, we applied two complementary mass spectrometry-based methods for the metabolomics analyses. Metabolite identification was confined to previously reported metabolites in spectral libraries, and therefore numerous detected metabolic features with interesting abundance patterns remain unidentified. In general, the correlation between mRNA and protein levels is only modest, which can be caused by e.g. variable protein turnover rates. Integration of protein turnover analysis to transcriptomics and proteomics was recently shown to increase the yield of identified disease gene candidates in pathological cardiac hypertrophy in mice,^52^ highlighting the benefit of adding further layers to omics-based analyses. Qualitative and quantitative assessment of all biological processes has been deemed essential for the investigation of cardiac metabolism by the American Heart Association.^53^ In the present study, integration of three omics analyses with four time points over the early postnatal period provides a comprehensive view to the metabolic changes taking place in the developing heart. Interestingly, in some cases we observed surprisingly long delays in the abundance changes from mRNA to protein and from protein to metabolite, as exemplified by the mevalonate pathway and ketogenesis. This highlights how crucial it is not only to use several omics methods but also to include several time points when analyzing dynamic phenomena with integrative systems biology approaches.

Considering the multicellular composition of the heart, the present study cannot elucidate cardiomyocyte-specific phenomena, as the data represent tissue-level changes. In cell volume, cardiomyocytes build up 70-80% of the cardiac tissue,^54^ whereas in cell numbers, cardiomyocytes constitute roughly one third of the cell population, while endothelial cells represent the most abundant cell type, with a ~45% proportion.^55^ These two cell types play a crucial role in post-infarction cardiac regeneration, and - unlike fibroblasts or leukocytes - fail to initiate the neonatal-type gene expression response upon injury in adult hearts.^26^ In cardiomyocytes, this was suggested to result from chromatin inaccessibility and thus represent an epigenetic roadblock that prevents cardiomyocyte cell cycle re-entry. Recent evidence suggests that metabolic pathways are intimately connected with epigenetic regulation.^56, 57^ Whether the postnatal metabolic remodeling is in fact the driving force that induces cardiomyocyte cell cycle exit requires further investigation.

In summary, we used an integrative systems biology approach with three levels of omics analyses in order to characterize changes in molecule abundances over the early postnatal period and to identify metabolic pathways associated with cardiac regeneration. To our knowledge, this is the first study combining transcriptomics with untargeted proteomics and global metabolomics analyses over several time points in the early postnatal heart, and as such provides an extensive resource of molecule abundances for future mechanistic studies. Furthermore, we present several examples of metabolic pathways with correlative changes on all omics levels. These include the well-established metabolic switch from glycolysis to fatty acid β-oxidation, but also many previously unreported changes in cardiac metabolic pathways, such as mevalonate pathway and ketogenesis. Finally, we identified a biological function for mevalonate and ketone body metabolism in the heart with a potential role in the regulation of neonatal cardiomyocyte proliferation. This integrated molecule-level data may open up new possibilities for the development of regenerative therapies.

## Acknowledgements

We thank Ms Marjo Vaha for technical assistance. RNAseq was carried out at Biomedicum Functional Genomics Unit (University of Helsinki, Finland) and high-content analysis using instrumentation at Biomedicum Imaging Unit (University of Helsinki, Finland).

## Sources of Funding

This study was supported by Business Finland (Tekes; project no. 40395/13, 3iRegeneration); the
Finnish Foundation for Cardiovascular Research; the Sigrid Jusélius Foundation; and the Academy of Finland (project no. 2666621).

## Disclosures

None.

